# The nucleotide exchange factor, GrpE, modulates substrate affinity by interaction of its N-terminal tails with the DnaK substrate-binding domain

**DOI:** 10.1101/2025.10.21.683677

**Authors:** Akshitha Maqtedar, Maria-Agustina Rossi, Eugenia M. Clerico, Robert V. Williams, Lila M. Gierasch

**Author notes:** For correspondence: Lila M. Gierasch. These authors contributed equally to this work (Akshitha Maqtedar, Maria-Agustina Rossi). Maria-Agustina Rossi: Instituto de Química Física Blas Cabrera (IQF), CSIC, E-28006 Madrid, Spain.

## Abstract

The 70-kDa heat shock proteins (Hsp70s) assist in protein folding through allosteric communication between their nucleotide-binding domains (NBDs) and substrate-binding domains (SBDs), which are connected by an interdomain linker. Their nucleotide-dependent allosteric cycle is modulated by ligand binding and co-chaperones, including nucleotide exchange factors (NEFs). GrpE, the NEF for the *E. coli* Hsp70, DnaK, has been proposed to have a dual effect on the chaperone, facilitating the exchange of ADP for ATP in the NBD in a temperature-dependent fashion and promoting substrate release from the SBD. We recently reported NMR-based evidence that GrpE binding to DnaK has a direct structural effect on the SBD. Here, we expanded on these findings and obtained new evidence for a model in which the disordered N-terminal tails of GrpE facilitate peptide dissociation from the nucleotide-free DnaK/GrpE complex by transiently binding to the canonical substrate-binding site in the SBD. This GrpE/SBD interaction, while weak, is favored by the high local concentration of the tails around the SBD after complex formation. Moreover, we identified the DnaK binding motif in GrpE’s N-terminal disordered tails as ^17^IIM^19^, which is conserved across many bacterial species. Excitingly, our data further suggest a mechanism for the temperature-dependence of GrpE’s modulation of DnaK’s refolding activity: as the temperature increases, unfolding of GrpE’s coiled-coil weakens its contacts with the SBD, reducing N-terminal tail binding, and thus increasing DnaK affinity to substrates.

## Introduction

Molecular chaperones play key roles in maintaining cellular protein health in all organisms (1). Central players among molecular chaperones are the 70-kDa heat shock proteins (Hsp70s) (2, 3). The structure of Hsp70s is conserved across prokaryotes and eukaryotes and consists of two domains–an N-terminal 44-kDa nucleotide-binding domain (NBD) and a C-terminal 30-kDa substrate-binding domain (SBD)–connected by a conserved interdomain linker. Hsp70s function *via* a nucleotide-dependent allosteric cycle based on a transition between two conformations, ATP- and ADP-bound, modulated by binding to incompletely folded substrate proteins and interactions with co-chaperones. Nucleotide-exchange factors (NEFs) facilitate the exchange of ADP for ATP in the NBD of all Hsp70s (3, 4). While the major role of NEFs is nucleotide exchange, it has also been observed that NEFs promote substrate release from the Hsp70 SBD. This functional role arises in part from NEF facilitation of the transition of Hsp70s from the ADP-bound high substrate affinity state to the ATP-bound low substrate affinity state, but may also be a consequence of a direct action of the NEF on the Hsp70 SBD (5–9).

Hsp70 NEFs are categorized into four classes: BAG proteins, HspBP1 and Hsp110 in eukaryotes, and GrpE in prokaryotes, mitochondria, and chloroplasts (3, 10). GrpE is the NEF of the *Escherichia coli* Hsp70 DnaK and is characterized by a distinctive structure and function. It is a homodimer with a unique cruciform shape consisting of a globular C-terminal region with two β-bundles and four α-helices, a long coiled-coil, and two N-terminal disordered tails (**Figure 1A**) (7, 11). The coiled-coil, characterized by a melting transition with a T^m^ of 48 °C, confers upon GrpE the ability to act as a cellular thermosensor (12, 13). At temperatures below 37 °C, GrpE assists in DnaK refolding activity of luciferase (14, 15). As the temperature increases, the GrpE coiled-coil domain starts to unfold, and GrpE loses its ability to assist DnaK in protein refolding (13, 16).

**Figure 1.**
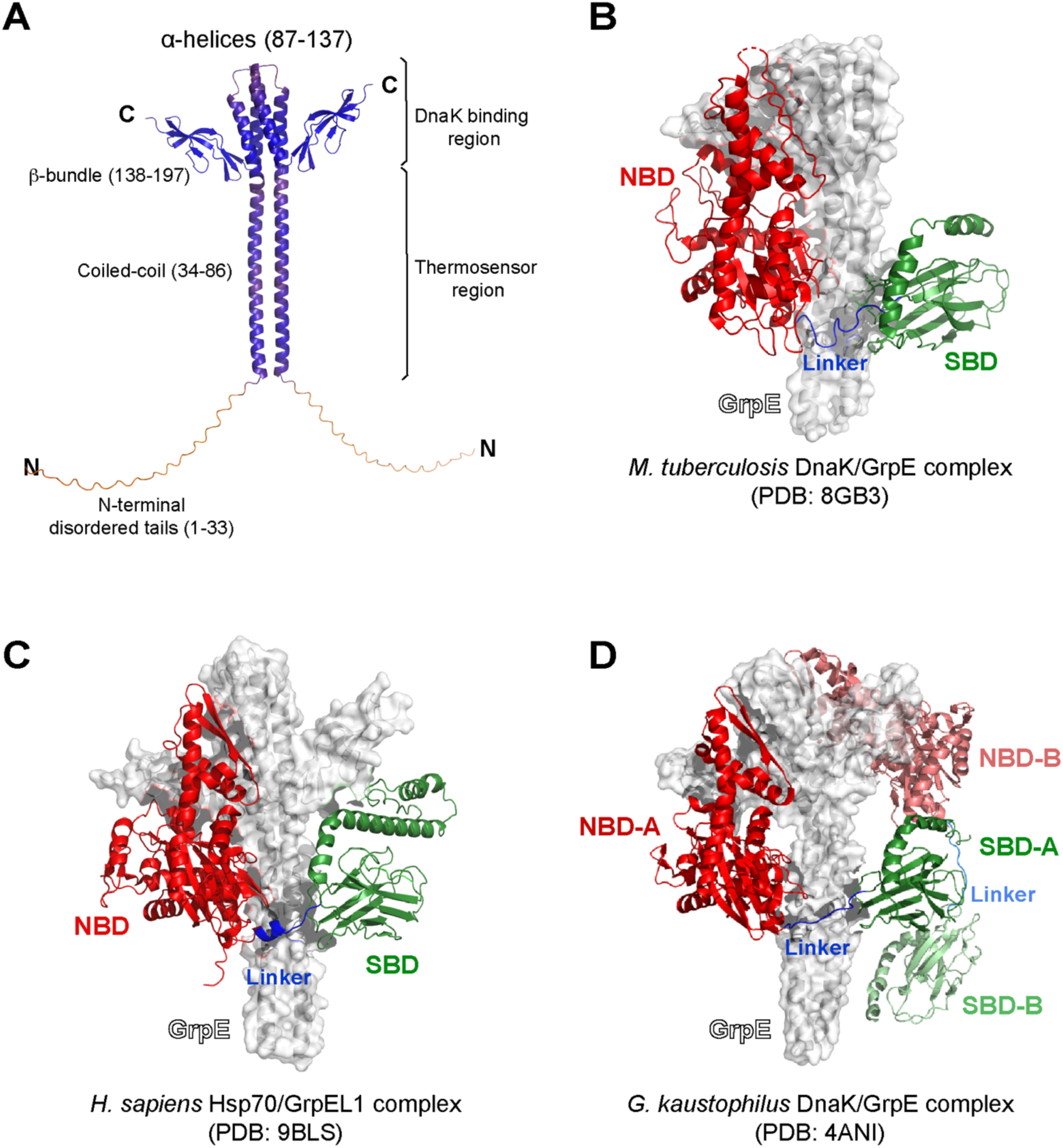
Structure of *E. coli* GrpE and DnaK/GrpE complexes of homologues to that of *E. coli*. *A,* AlphaFold-based structure of *E. coli* GrpE where the structural regions are indicated and colored by AlphaFold confidence (red = low confidence, blue = high confidence). *B-D*, The structures of diverse complexes between Hsp70s and GrpE variants from various species are shown. GrpE is always highlighted in white, and in the Hsp70, the NBD is in red, the SBD is in green, and the interdomain linker is in blue. *B,* CryoEM structure of the DnaK/GrpE complex from *Mycobacterium tuberculosis* (PDB: 8GB3); *C,* cryoEM structure of the mitochondrial GrpE/Hsp70 complex from *Homo sapiens* (GrpEL1 and Mortalin ; PDB: 9BLS); *D,* crystal structure of the DnaK/GrpE complex from *Geobacillus kaustophilus* (PDB: 4ANI), in which we show one DnaK bound to GrpE (NBD-B: salmon; SBD-B: light green; interdomain linker: sky blue).

GrpE appears to exploit the dual role described above for NEFs in general in its facilitation of the role of DnaK in protein refolding. First, GrpE accelerates the Hsp70 allosteric cycle *via* its well-established promotion of exchange of ADP by ATP. For this role, GrpE binds to the ADP-bound NBD and increases the nucleotide off-rate by stabilizing subdomain IIB in lobe II in a conformation that opens the nucleotide-binding cleft (4, 10). In this mechanism, it is important to note, as we reported previously (17), that the ADP-bound NBD/GrpE complex is of low stability and has increased conformational heterogeneity in comparison to the stable nucleotide-free NBD/GrpE complex. Thus, GrpE promotes the release of ADP, poising the NBD for ATP binding, and facilitating the resulting conformational shift of DnaK to a low substrate affinity state. The second role of GrpE has been suggested by several studies that report weakening of the binding affinity of substrate to DnaK upon binding to GrpE (5–7). In an early structural study of the GrpE complex with DnaK, Harrison *et al.* reported that full-length GrpE weakened the binding of DnaK to an unfolded polypeptide substrate, but that N-terminally truncated GrpE did not (7). Follow-up work has explored how the kinetics of substrate binding to DnaK are affected by binding of GrpE (5, 6, 9) and the necessity for the 33-residue N-terminal mobile region to be present for the enhancement of substrate release (5–7, 9).

In terms of a mechanism for the impact of GrpE binding on DnaK substrate affinity, previous studies raised the possibility that the 33-amino acid N-terminal disordered tails interact with the canonical substrate-binding site in the DnaK SBD to assist in peptide release (5–9), but no direct interaction has been observed thus far. Removal of the N-terminal 33 amino acids (in a construct called GrpE^33-197^) abrogated the impact of GrpE complex formation on substrate binding (7, 9). Notably, the previous studies examining these effects of DnaK/GrpE complex formation on substrate affinity focused on ADP-bound DnaK, which our previous study suggested is not the favored species in the ensemble of GrpE-bound DnaK chaperones. In previous and current studies, we have examined the complex of GrpE and nucleotide-free DnaK, given that our observations supported the greater stability and conformational homogeneity of this state.

In addition, in our previous study, we reported NMR chemical shift perturbations (CSPs) in the DnaK SBD upon GrpE binding to DnaK, which supports the earlier biochemical findings of GrpE-induced altering of substrate affinity (5–7, 9). Two possible mechanisms for the impact of GrpE binding on the SBD were suggested: (1) transmission of an allosteric signal from the NBD to the interdomain linker, and then to the SBD or (2) a direct interaction between the GrpE N-terminal disordered tails and the DnaK SBD (17).

There are now three atomic resolution structures of DnaK/GrpE complexes from *E. coli* DnaK homologues (8, 18, 19) that provide structural insights into the nature of the interaction between *E. coli* GrpE and the DnaK SBD: the cryo-EM structure of the human Mortalin/GrpEL1 complex (**Figure 1C**) (18), the x-ray crystal structure of the *G. kaustophilus* DnaK/GrpE (**Figure 1D**) (19), and the cryo-EM structure of the *M. tuberculosis* DnaK/GrpE complex (**Figure 1B**) (8). These structures reveal three significant contact regions between the NEF and Hsp70, which are designated I to III using descriptors from the structure of the *M. tuberculosis* GrpE/DnaK complex (**Figure 1B**) (8). Contact regions I and II involve interactions between the NEF and the NBD. Most exciting for the possible action of the NEF on substrate binding is the location of contact region III, wherein the DnaK SBD interacts with the coiled-coil domain of GrpE. Here, DnaK SBD loop R388-P393 (loop 1) interacts with the D68-R75 segment of the coiled-coil domain of GrpE, and DnaK residue D451 interacts with R64 from GrpE (*M. tuberculosis* numbering). In addition, the interdomain linker lies across the GrpE coiled-coil domain. Comparable interactions are observed, albeit with lower resolution, in the other two GrpE/DnaK structures of *E. coli* DnaK homologues (18, 19). While all of these structures provide provocative information on the interaction of the coiled-coil region of GrpE with the DnaK interdomain linker and SBD, in none of them are the disordered N-terminal tails resolved, leaving their potential role in mediating substrate release unclear.

Here, to explore the impact of *E. coli* GrpE binding on the substrate affinity of DnaK, we have deployed a multidisciplinary approach. First, we use current powerful prediction methods to support the structural homology of the *E. coli* GrpE/DnaK complex with those of the homologues. Importantly, we find that key interaction regions in all the complexes are highly conserved across a broader set of GrpE/DnaK homologues, supporting the generality of the structure of the complex. We next explored in greater depth how the dynamic N-terminal tails of GrpE may participate in the mechanism of GrpE action on DnaK. Here we rely heavily on NMR spectroscopy, as it is well suited to studying highly dynamic regions and can shed light on the involvement of the GrpE N-terminal tails in the complex. On the basis of our findings, we propose a model in which the disordered N-terminal tails of GrpE actively facilitate peptide dissociation from the apo-DnaK/GrpE complex by competing with the exogenous substrate for binding to the SBD canonical substrate-binding site. Our data argue that the N-terminal disordered tails transiently bind to the SBD substrate-binding site with low affinity. Despite the low affinity of the N-terminal tail binding interaction, the covalent linkage of the tails to GrpE in the complex confers upon them a high local effective concentration. Furthermore, we have identified the DnaK binding motif in the GrpE N-terminal tail as ^17^IIM^19^, which is conserved across many bacterial species. Finally, our model provides an explanation for the reduced impact of GrpE/DnaK binding on substrate affinity and the loss of DnaK refolding activity at elevated temperatures (14, 15, 20). As temperature increases, the coiled-coil domain of GrpE begins to unfold (13, 16), which weakens the GrpE contacts with the SBD, reducing the likelihood of the direct interaction of the N-terminal tails with the SBD and thus abrogating the GrpE effect on peptide release.

## Results

### Comparison of the predicted structure of the E. coli DnaK/GrpE complex and DnaK/GrpE complexes from homologues supports structural conservation

As there has not been a structure experimentally determined structure for the complex of *E. coli* GrpE and full-length DnaK, we used AlphaFold2 to predict this structure (21, 22) (**Figure 2A**). As seen in the previously reported structures of complexes from homologous organisms, the *E. coli* DnaK NBD and SBD in the complex with GrpE are not associated with one another, similar to the ADP-bound state of *E. coli* DnaK (23). The NBD/GrpE interactions are consistent with the previous x-ray structure of GrpE complexed with the isolated *E. coli* DnaK NBD (7) and the corresponding predicted structure from our lab (17). Excitingly, the contacts formed between the GrpE coiled-coil with DnaK are the same in the *E. coli* complex as in those of homologues (8, 18, 19). These include interaction of the coiled coil with SBD loop K414-P419 (loop 1), SBD residues N415, T417, D477, and the interdomain linker (*E. coli* numbering). The involvement of the interdomain linker in binding to the coiled-coil of GrpE is also consistent with previous biochemical observations showing a loss of proteolytic lability of this region in the DnaK/GrpE complex relative to its susceptibility in DnaK (6). Thus, three major interactions are implicated in the DnaK/GrpE complex: the DnaK NBD with the GrpE head, the DnaK interdomain linker with the GrpE coiled-coil, and SBD loop 1 and residues N415, T417, and D477 with the GrpE coiled-coil.

**Figure 2.**
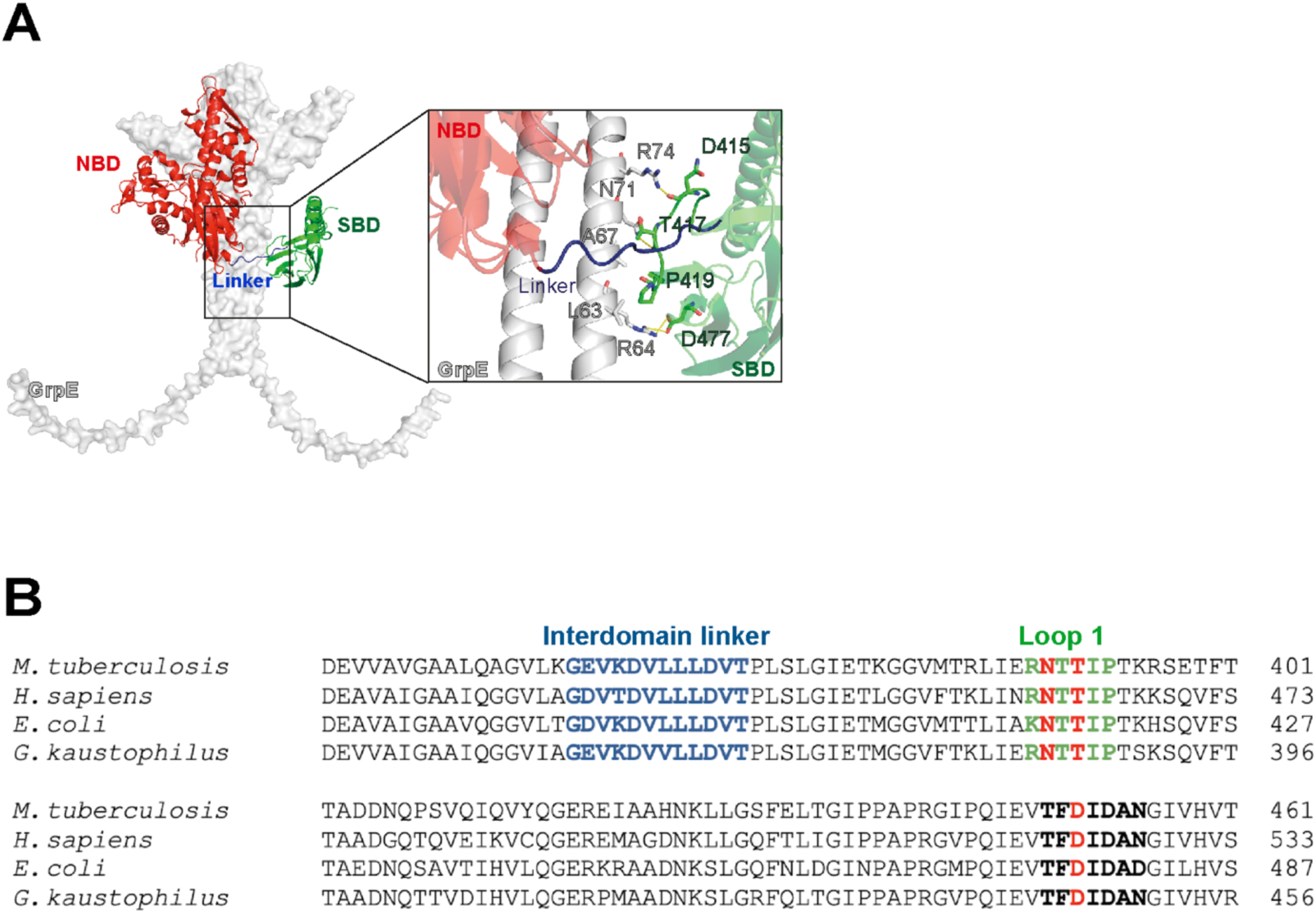
Comparison of the AlphaFold2-predicted *E. coli* DnaK/GrpE complex to DnaK/GrpE complexes from other species reveals conserved interaction modes. *A,* AlphaFold2-predicted structure of the *E. coli* DnaK/GrpE complex and a close-up view of the interaction interface between DnaK SBD and GrpE coiled-coil domain. GrpE is highlighted in white, and in the Hsp70, the NBD is in red, the SBD is in green, and the interdomain linker is in blue. The interacting residues in GrpE are shown in light grey and in DnaK are shown in green. *B,* Sequence alignments of DnaK from *M. tuberculosis,* human (*mortalin,* HSPA9)*, E. coli,* and *G. kaustophilus*. The interdomain linker sequence is highlighted in blue. Loop 1 is in green (*E. coli* K414-P419, *G. kaustophilus* R382-P388, *H. sapiens* Mortalin HSPA9 R460-P465, and *M. tuberculosis* R388-P393). The residues that contact the GrpE coiled-coil domain are highlighted in red: Asn (*E. coli* N415, *G. kaustophilus* N383, Mortalin HSPA9 N461, and *M. tuberculosis* N389), Thr (*E. coli* T417, *G. kaustophilus* T385, Mortalin HSPA9 T463, and *M. tuberculosis* N391) and Asp (*E. coli* D477, *G. kaustophilus* D446, Mortalin HSPA9 D523, and *M. tuberculosis* D451).

This comparison of the predicted *E. coli* DnaK/GrpE complex with structures of homologue complexes leads to the expectation that the interaction interfaces are conserved across species. An alignment of DnaK sequences from *E. coli*, *G. kaustophilus*, *H. sapiens* mitochondrial, and *M. tuberculosis* reveals sequence identities ranging from 54 to 59% and similarities between 70 and 74%. Notably, the interdomain linker is highly conserved across all four variants, with nearly identical residue sequence and length (**Figure 2B**). Loop 1 also maintains a conserved residue sequence, particularly the SBD N415 and T417 residues that interact with GrpE, which are preserved in all four species (**Figure 2B**). Additionally, the SBD D477 residue and the surrounding region exhibit identical sequence identity across the four Hsp70s analyzed (*E. coli* numbering) (**Figure 2B**).

We also explored the conservation of the structural features in the complexes with DnaK among different GrpEs. Representative GrpEs from different organisms were included in an alignment: GrpEL1 from *H. sapiens* mitochondria (eukaryotes), GrpE from *E. coli* and *K. pneumoniae* (gram-negative bacteria), and GrpE from *M. tuberculosis*, *G. kaustophilus*, and *B. subtilis* (gram-positive bacteria). GrpE homologues from gram-negative species show exceptionally high overall sequence conservation, with 87% identity and 94% similarity, while the human and gram-positive counterparts exhibit lower overall identity and similarity percentages. However, all analyzed GrpEs display conserved stretches flanked by non-conserved regions (**Figure 3A and 3B**). In particular, the residue sequence of the interaction interfaces between GrpE and DnaK NBD are highly conserved: residues I91-D110 in the β-bundles, which interact with DnaK subdomain IB, and the stretch L148-I157 in the α-helices, which interacts with DnaK subdomain IIB (*E. coli* numbering) (**Figure 3B**). Notably, G122, which, when mutated to an Asp residue, disrupts the interaction between DnaK NBD and GrpE, and renders the NEF inactive (17, 24), is conserved across most variants. The motif interacting with DnaK SBD in the coiled-coil domain (residues L63-R75, *E. coli* numbering) also shows high sequence identity.

**Figure 3.**
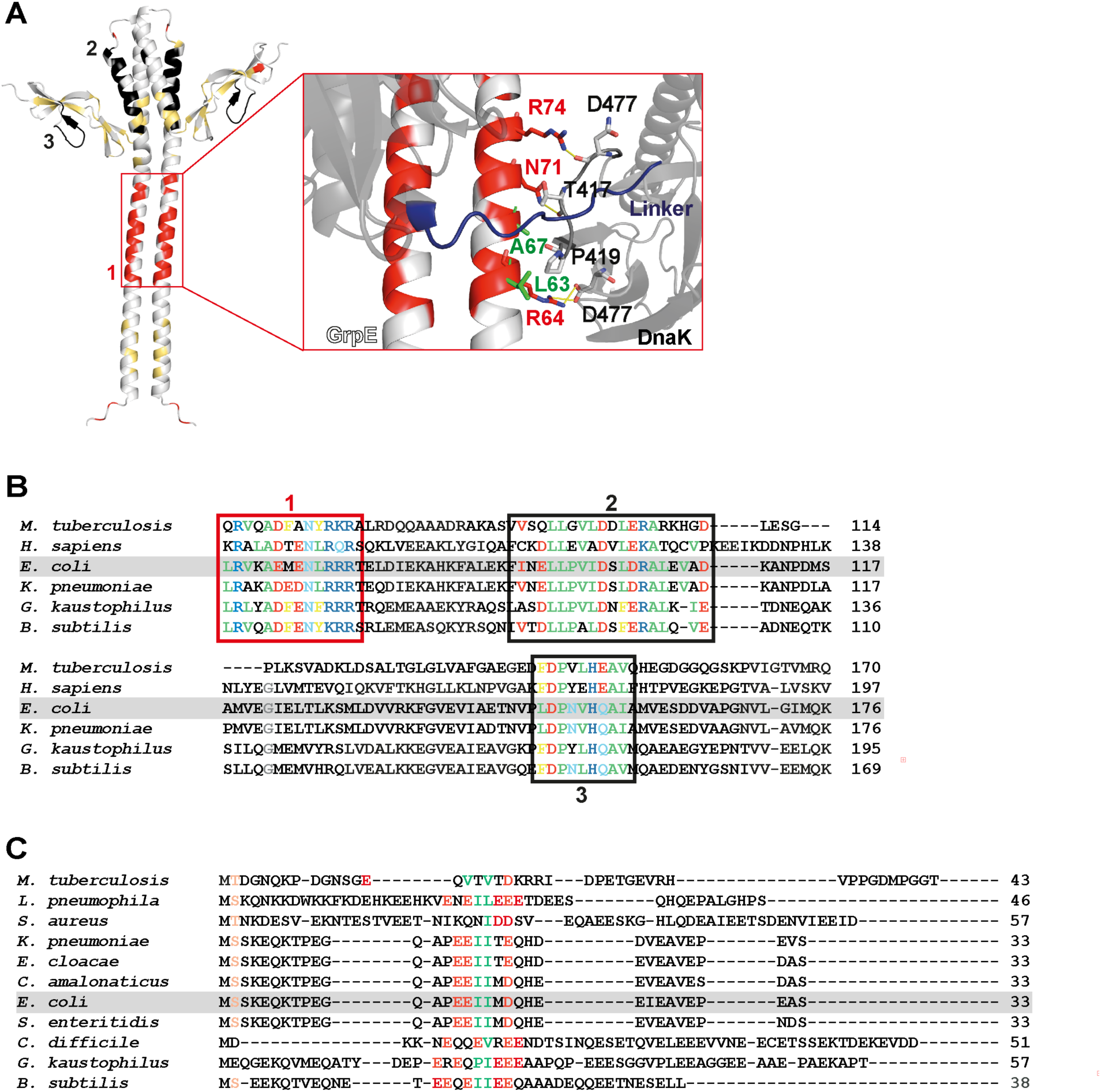
GrpE homologues exhibit sequence conservation in the interaction interfaces of the DnaK/GrpE complex. *A,* Conserved residues are mapped in the *E. coli* GrpE structure: (1) L63-R75 in red, (2) I91-D110 in black, and (3) L148-I157 in black. All other conserved residues outside these areas are in yellow. The zoomed in region shows the interaction interface between the GrpE coiled-coil domain and DnaK SBD. *B,* Sequence alignment of GrpE homologues where conserved residues are highlighted (hydrophobic in green, polar in light blue, negatively charged in red, positively charged in blue, aromatic in yellow, and G in grey). The (1) L63-R75 region in the coiled-coil domain that interacts with DnaK is highlighted with a red rectangle; The (2) I91-D110 region that contacts DnaK subdomain IB and the (3) L148-I157 region that interacts with DnaK subdomain IIB are highlighted with a black rectangle. The numbering in this caption corresponds to the *E. coli* GrpE. *C,* Sequence alignments for the N-terminal disordered tails of GrpE. The only sequence similarity in the N-terminal tails is highlighted (hydrophobic residues in green, negatively changed ones in red).

Given the proposed role of the N-terminal disordered tails of the *E. coli* GrpE in modulating the affinity of DnaK for a model peptide (5–8), we examined sequence similarities for this region as well in the alignments (**Figure 3C**). The N-terminal disordered tails of GrpE exhibit minimal sequence similarity among the set of structurally described homologues. To explore any potential conservation that did not emerge in this small set of sequences, we included additional prokaryotic homologues in our alignment, and two key features were observed: a high proportion of negatively charged residues, primarily Glu, and a conserved pattern of two or three hydrophobic residues flanked by negatively charged amino acids, typically Glu. In *E. coli*, these hydrophobic residues correspond to I17, I18, and M19.

### Binding of GrpE reduces the substrate-binding affinity of nucleotide-free E. coli DnaK, but binding of GrpE lacking its N-terminal 33-amino acids does not

Previous studies reported that GrpE complex formation with DnaK influences the kinetics of peptide binding and release in the presence of ADP (5–7, 9). Given that the nucleotide-free state is the functionally critical state of the DnaK NBD complex with GrpE (17), we tested the impact of GrpE binding to apo-DnaK on substrate binding. In this experiment, the amount of DnaK relative to peptide and GrpE was varied. We found that the apparent affinity (app-*K*^D^) of DnaK for the fluorescein isothiocyanate (FITC)-labeled peptide p5 (ALLLSAPRR, (25)) was reduced in the presence of GrpE (**Table 1**, **Figure 4A and S5A**). Importantly, removal of the N-terminal disordered tails (in GrpE^33-197^) abolished this effect such that the observed substrate affinity was essentially the same as that of isolated DnaK. We also found that in the presence of increasing concentrations of NEF, GrpE^33-197^ is still unable to stimulate peptide release (**Figure 4B**). Thus, our results using apo-DnaK support a role for the GrpE tails in modulating DnaK substrate affinity, as previously described in the case of ADP-bound DnaK (5–7, 9).

**Figure 4.**
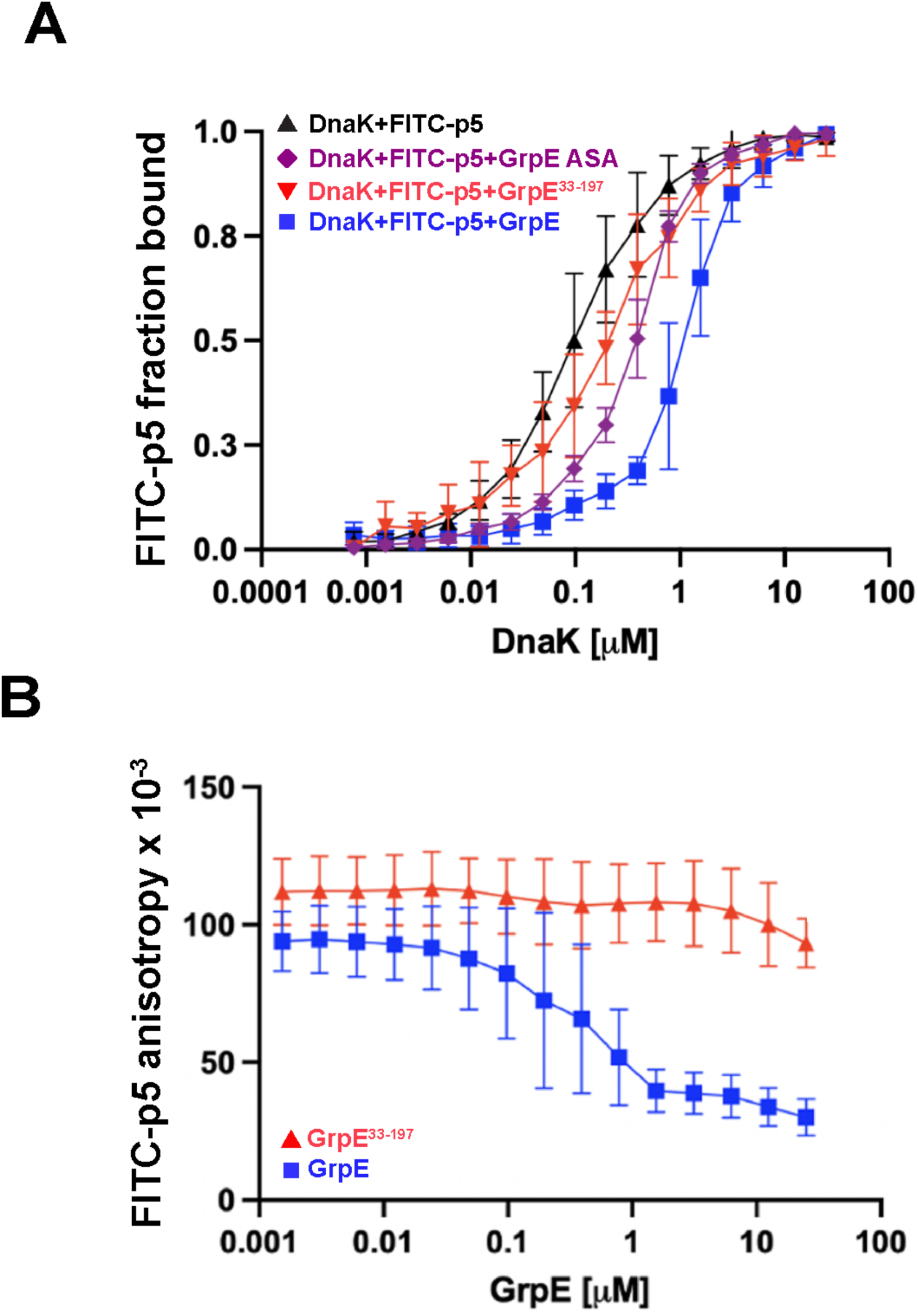
Binding of GrpE reduces affinity of the DnaK SBD for a model peptide. *A,* Apparent affinity of DnaK for the model peptide FITC-p5 in the absence or presence of 1 μM GrpE, GrpE^33-197^, and GrpE ASA at 22 °C. *B,* Apparent affinity of DnaK for the model peptide FITC-p5 when the concentration of GrpE and GrpE^33-197^ was varied at 22 °C.

### NMR chemical shifts observed for the DnaK complex with GrpE argue that an Ile residue occupies the substrate-binding groove of the SBD

To further investigate the impact of GrpE binding on the SBD of DnaK, we collected HMQC NMR spectra of nucleotide-free,^13^C-methyl ILV-DnaK in complex with GrpE or GrpE^33-197^. When DnaK forms a complex with GrpE, there are many chemical shift changes relative to free DnaK, including small and localized changes associated with the SBD. By contrast, formation of the DnaK complex with GrpE^33-197^ results in fewer perturbations within the SBD, supporting a role for the disordered N-terminal tails (**Figure S1**). We focused our NMR analysis of the GrpE complex using δ1-methyl chemical shifts of Ile 401 and 438 of the DnaK SBD, as they have proven to be extremely useful reporters of residue binding to the canonical substrate-binding pocket of the SBD (26, 27). In the DnaK/GrpE complex, I438 exhibited a chemical shift indicative of binding of an Ile residue to the central pocket of the substrate-binding cleft (**Figure 5A**) (27). This peak shift is only observed in the DnaK/GrpE complex and not in nucleotide-free DnaK or in DnaK in complex with GrpE^33-197^ (**Figure 5B**), supporting occupancy of the SBD binding cleft by the GrpE N-terminal disordered tails. GrpE has two Ile in its tail (sequence ^14^PEE**II**MDQH^22^), and both are conserved among the different GrpE homologues (**Figure 3C**). It is not possible to speculate about which Ile in the N-terminal tail is bound to the SBD pocket; however, the broad and low intensity of the SBD I438 δ1-methyl resonance suggests that the binding affinity of the GrpE N-terminal tail for the SBD is low, fully consistent with the numerous negative charges in the region of the Ile residues that may be binding to the SBD.

**Figure 5.**
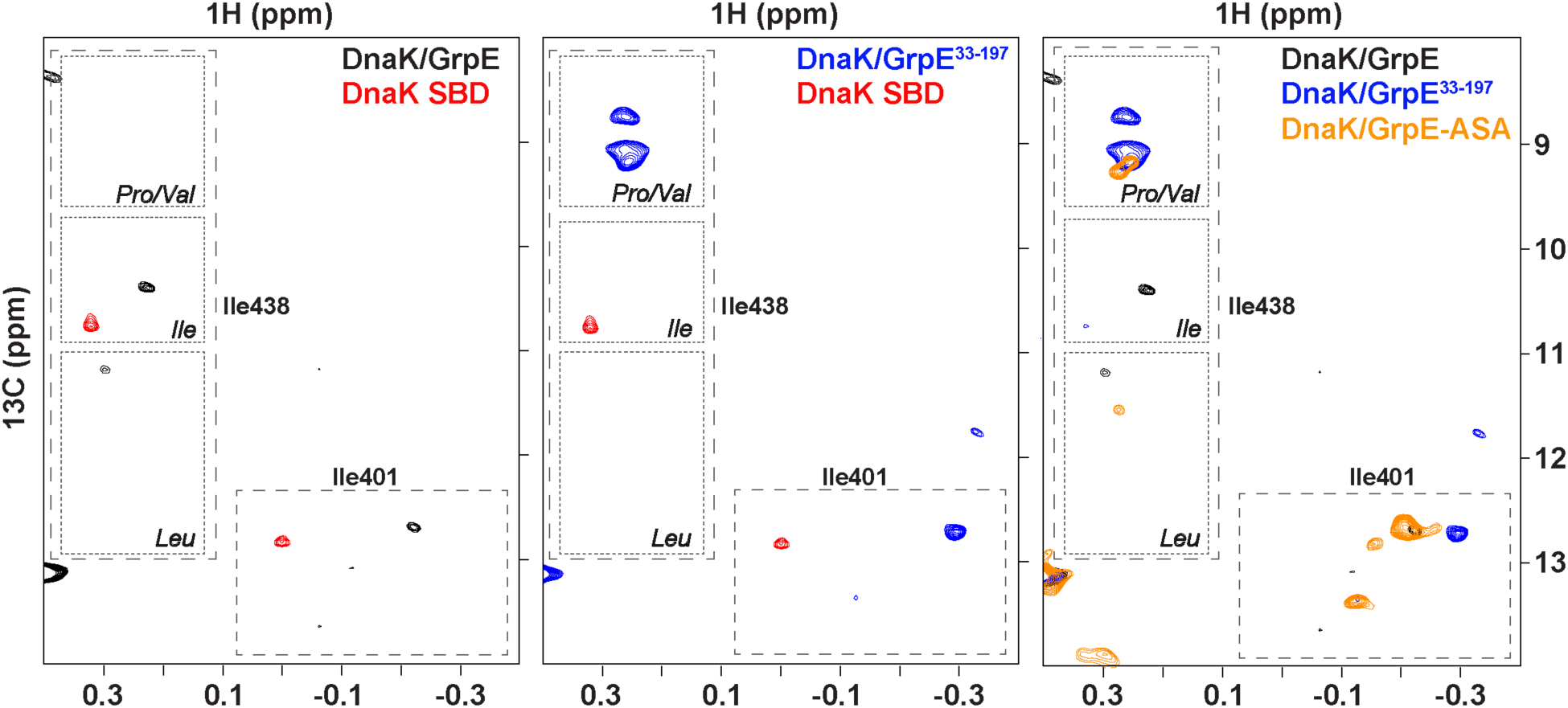
DnaK SBD reporter NMR signals, I401 and I438, indicate which residues bind to the central pocket when complexed with GrpE and its variants. Ile resonances of the substrate binding pocket on the HMQC at 25 °C of (Left) ^13^C-methyl labeled ILV-DnaK/GrpE complex (black) and isolated apo-DnaK SBD (red), (Middle) ^13^C-methyl labeled ILV-DnaK/GrpE^33-197^ complex (blue) and isolated apo-DnaK SBD (red), and (Right) ^13^C-methyl labeled ILV-DnaK/GrpE complex (black), ^13^C-methyl labeled ILV-DnaK/GrpE^33-197^ complex (blue), and ^13^C-methyl labeled ILV-DnaK/GrpE ASA complex (orange).

Interestingly, in the DnaK/GrpE^33-197^ complex, the position of the I438 resonance is consistent with a Val or Pro residue occupying the central pocket (**Figure 5B**) (27). While we lack a definitive explanation for this observation, we offer two potential explanations: One possibility is that a region of GrpE other than the N-terminal tail is occupying the canonical substrate-binding site in this complex. This idea is supported by the fact that the Pro/Val peak in the spectrum of the DnaK/GrpE^33-197^ complex disappears upon addition of the p5 peptide **(Figure S2)**, suggesting that the Pro/Val peak seen in the DnaK/GrpE^33-197^ complex indeed arises from interactions at the canonical substrate-binding site. Alternatively, the binding of GrpE that lacks the N-terminal tails may induce a conformational change in the SBD. Since the DnaK/GrpE^33-197^ complex shows substrate-binding affinity similar to that of nucleotide-free DnaK, it is possible that the α-helical lid adopts a more closed conformation over the β-subdomain of the SBD in the complex with GrpE^33-197^. This conformational state may be characterized by a shift in the resonance of I438. Further studies will be required to distinguish between these or any other possibilities.

### NMR analysis supports binding of GrpE N-terminal tails to the canonical substrate-binding site in the DnaK SBD through the ^17^IIM^19^ motif

Our NMR analysis shows that upon GrpE binding the Ile δ1-methyl chemical shift of DnaK I438 is consistent with an Ile residue occupying the central pocket. We thus hypothesized that the conserved ^17^IIM^19^ motif in the N-terminal tails binds to the canonical substrate-binding site in the DnaK SBD and that its binding competes with substrate binding thereby facilitating substrate release. To test this model, we mutated I17, I18, and M19 to A17, S18, and A19; this GrpE variant is named GrpE ASA. The mutant sequence is significantly less hydrophobic than the native IIM, thus it is predicted to bind more weakly to the central binding pocket, which typically favors nonpolar residues (27). If the ASA sequence ablates the tail interaction with the SBD as we predict, GrpE ASA is not expected to greatly modulate the affinity of nucleotide-free DnaK for the model peptide FITC-p5. Indeed, binding of GrpE ASA did not decrease the affinity of DnaK for its substrate (**Table 1**, **Figure 4A and S5A**), as compared to GrpE, indicating that ^17^IIM^19^ substitution for ASA residues renders GrpE almost as ineffective as removal of the N-terminal tails. These results are consistent with our hypothesis that the ^17^IIM^19^ motif in the N-terminal disordered tails of GrpE binds to the canonical substrate-binding cleft of DnaK SBD.

We also analyzed the I438 region in the NMR spectrum of ^13^C-methyl ILV-labeled apo-DnaK in complex with GrpE ASA (**Figure 5C**) (27). In this complex, the peak corresponding to an Ile residue in the central pocket of the SBD observed in the apo-DnaK/GrpE spectrum disappeared, and a new peak emerged in the region assigned to I438 when a Pro/Val occupies the central substrate-binding site. This peak position is close to that seen in the DnaK/GrpE^33-197^ spectrum, suggesting that similar mechanisms may be responsible for these chemical shifts. Overall, the substitution of the conserved ^17^IIM^19^ motif by ASA leads to a loss of the ability of GrpE to stimulate substrate dissociation from DnaK’s central pocket, likely due to the absence of the binding motif.

### GrpE interactions with DnaK lead to both conformational and dynamic changes in GrpE N-terminal tails

By observing ^15^N-labeled GrpE as it forms a complex with unlabeled DnaK we were able to directly monitor chemical shift changes and alterations in dynamics of the NEF. Due to the size and anisotropy of the GrpE dimer, most of its NMR resonances are broadened and low intensity. However, the ^1^H-^15^N HSQC spectrum of full-length ^15^N-labeled GrpE reveals around 30 distinct peaks (**Figure 6A**), all of which are absent in the spectrum of ^15^N-labeled GrpE^33-197^ (**Figure S3**), leading to the tentative assignment of these resonances to the N-terminal 33 residues. Assignment of this sequence, carried out following standard procedures (see Methods), confirmed that these resonances are from residues in the flexible N-terminal tails. Moreover, the observation of relatively narrow resonances for the N-terminal tails is fully consistent with the mobility of this region and the inability of either x-ray crystallography or cryo-EM methods to observe the N-terminal tails. For our study, the fact that this region is NMR observable is a highly informative finding.

**Figure 6.**
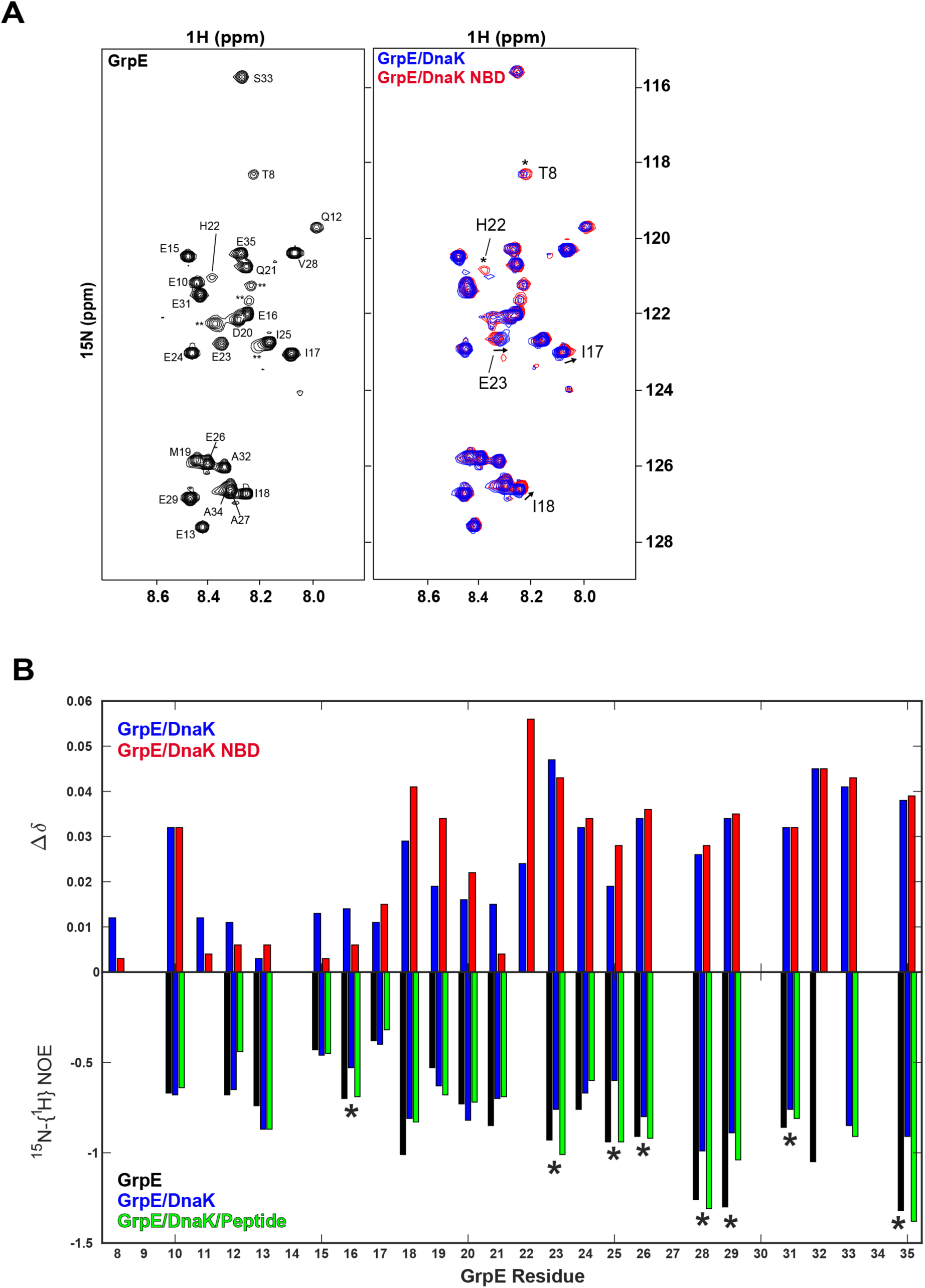
^15^N-GrpE NMR experiments support transient binding of GrpE N-terminal tails to DnaK SBD. *A,* (Left) Assigned ^1^H-^15^N HSQC spectrum of the full-length ^15^N-labeled GrpE (black) and (Right) ^15^N-GrpE in complex with full-length DnaK (blue) or the DnaK NBD^1-392^ (red) showing the largest chemical shift perturbations (CSPs), observed for residues T8, I17, I18, H22, and E23. *B,* (Top) Bar plot comparing CSPs (Δδ) for ^15^N-GrpE in complex with full-length DnaK (blue) or the DnaK NBD^1-392^ (red) relative to ^15^N-GrpE alone. CSPs were not measured for proline residues (P9, P14, P30) or due to resonance overlap (A27, A34). (Bottom) ^15^N-{^1^H}-NOE values for GrpE (black), GrpE/DnaK (blue), and GrpE/DnaK/Ash2 peptide (green) complexes. Residues marked with asterisks are consistent with the tail binding hypothesis, as the ^15^N-{^1^H}-NOE decreases in presence of DnaK and comes back to levels similar to GrpE alone upon addition of the Ash2 peptide. Several NOEs were not measured due to a proline residue (P9, P14, P30) overlapping resonances (A27, A34), or insufficient intensity in the saturated spectrum (H22).

A comparison of the chemical shifts of ^15^N GrpE free and in complex with DnaK shows perturbations in residues I17, I18, H22, and E23 of the GrpE N-terminal tails. Importantly, these CSPs are only present when full length DnaK is used (**Figure 6B**). These shifts are consistent with our model that the GrpE N-terminal tails interact transiently with the DnaK SBD. Importantly, the perturbations observed in I17 and I18 support the direct involvement of the ^17^IIM^19^ motif in this interaction.

To further probe the dynamics of this interaction, we performed ^15^N heteronuclear NOE experiments on ^15^N-GrpE, the ^15^N-GrpE/DnaK complex, and ^15^N-GrpE/DnaK in the presence of high concentrations of a high-affinity substrate peptide (Ash2: CRLLPLLFTPSR), which occupies the DnaK substrate-binding site and out-competes weaker binding substrates (27). In isolation, the N-terminal residues of GrpE exhibited negative NOE values, consistent with a disordered region. Upon DnaK binding, a subset of residues, particularly those following the ^17^IIM^19^ motif, showed increased (less negative) NOE values, indicating reduced mobility, which would arise from transient engagement with the SBD (**Figure 6B**). Importantly, addition of the Ash2 peptide restored the NOE values of the residues nearby the ^17^IIM^19^ to similar levels as seen in the free state. This observation provides strong support for binding of the GrpE tails to the substrate-binding cleft of the SBD, such that peptide binding displaces the tails (**Figure 6B**). These results strongly support a model in which the GrpE N-terminal tails, *via* the ^17^IIM^19^ motif, transiently bind the canonical substrate-binding site of the DnaK SBD. This interaction reduces tail mobility and is sensitive to substrate competition, highlighting a regulatory mechanism for substrate release.

### The impact of the GrpE N-terminal tails on peptide binding to the DnaK SBD is temperature-dependent

*In vivo, E. coli* GrpE acts as a thermosensor for the DnaK allosteric cycle and modulates DnaK activity in response to temperature changes to maintain proteostasis (14, 15). Therefore, we tested whether the modulation of DnaK’s substrate affinity by GrpE is affected by temperature and how our proposed role for the GrpE N-terminal tails might inform the temperature dependent function of GrpE.

We first determined DnaK’s apparent affinity for the FITC-p5 peptide (app *K*_D_) at 38 °C, a temperature reported to abrogate the NEF activity of GrpE (13, 16) in the absence or presence of GrpE or GrpE^33-197^ (**Table 1**, **Figure 7A and S5B**). At 38 °C, the apparent affinity of DnaK for FITC-p5 is essentially unchanged upon addition of either GrpE or GrpE^33-197^, in contrast to the reduction of app-*K*_D_ of DnaK for this peptide upon binding GrpE at 22 °C. To explain this finding, we hypothesize that the temperature-induced unraveling of GrpE’s coiled-coil domain (14) disfavors the transient interaction of the GrpE N-terminal tails with the substrate-binding pocket of DnaK SBD.

**Figure 7.**
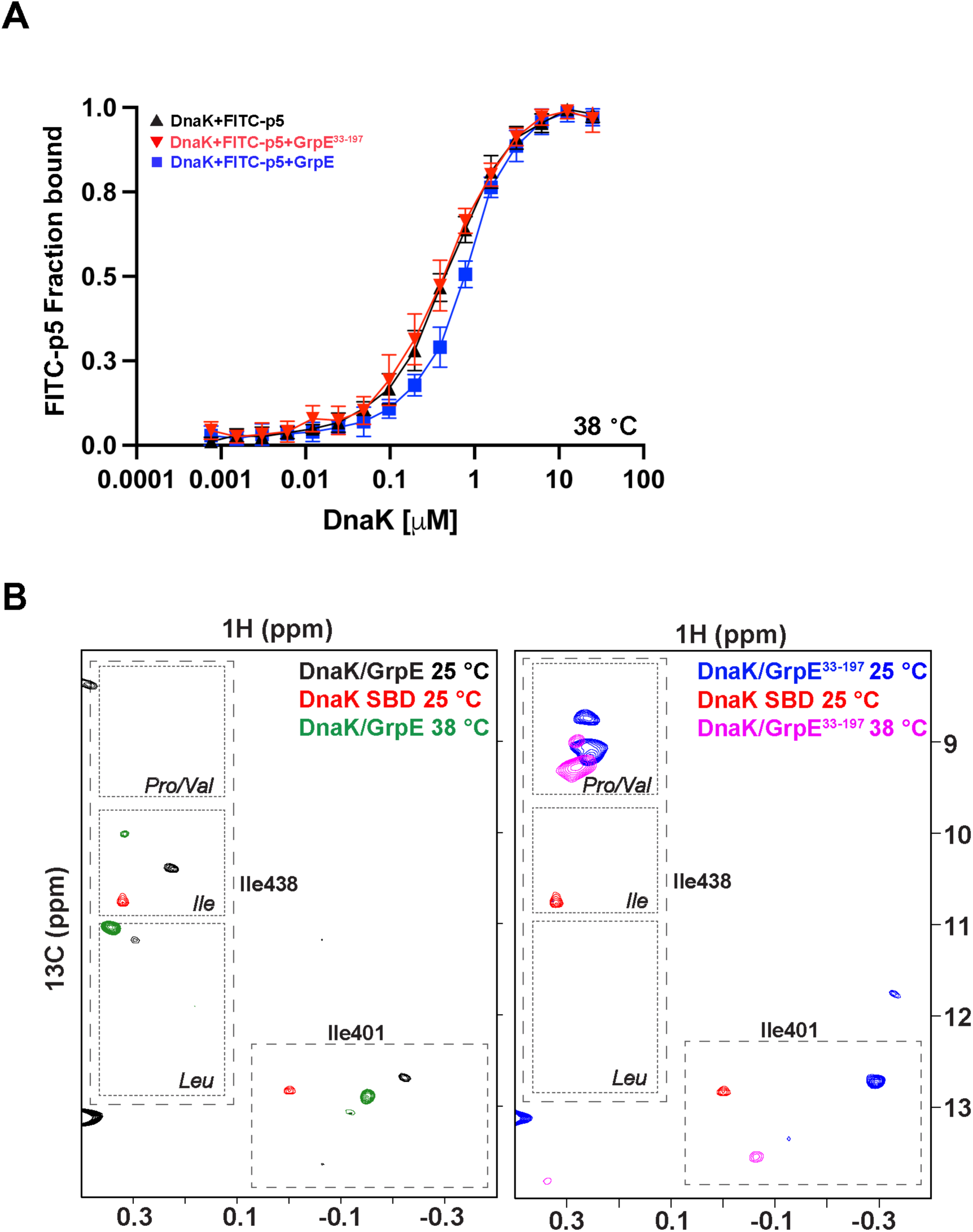
GrpE modulates peptide release from apo-DnaK less at elevated temperatures. *A,* Apparent affinity of DnaK for the model peptide FITC-p5 in the absence or presence of 1 μM GrpE and GrpE^33-197^ at 38 °C. *B,* Ile resonances of the substrate binding pocket on the HMQC of (Left) ^13^C-methyl labeled ILV-DnaK/GrpE complex (black) at 25 °C, ^13^C-methyl labeled ILV-DnaK/GrpE complex (green) at 38 °C, and isolated apo-DnaK SBD (red) at 25 °C and (Right) ^13^C-methyl labeled ILV-DnaK/GrpE^33-197^ complex (blue) at 25 °C, ^13^C-methyl labeled ILV-DnaK/GrpE^33-197^ complex (pink) at 38 °C, and isolated apo-DnaK SBD (red) at 25 °C.

To test this hypothesis, the HMQCs of nucleotide-free ^13^C-methyl labeled ILV-DnaK bound to GrpE or GrpE^33-197^ were obtained at 38 °C (**Figure 7B**). At 38 °C, the SBD I438 resonance in the DnaK/GrpE complex disappears from the region that corresponds to occupation of the substrate-binding pocket by an Ile residue, as was observed at 25 °C. In contrast, one new I438 resonance appears. The position of the one I438 resonance corresponds with the unbound conformation of the isolated DnaK SBD. Interestingly, the SBD I438 resonance position for the DnaK/GrpE^33-197^ complex is at the same position regardless of the temperature (25 or 38 °C), and as described above, this position is compatible with a Pro or Val in the central pocket.

These experiments suggest that DnaK is able to bind the GrpE N-terminal tails at the temperature when GrpE retains its NEF activity and modulates substrate affinity, but when the temperature is raised such that the coiled-coil domain begins to unfold (13, 16), the interaction of the ^17^IIM^19^ motif in the GrpE N-terminal tails and the substrate-binding pocket is disfavored. Overall, these results help to explain why GrpE cannot assist DnaK refolding activity at higher temperatures.

## Discussion

Nucleotide exchange is pivotal to physiological modulation of the allosteric cycle of Hsp70s. NEFs play a key role in this process, operating through a common mechanism. All NEFs bind to the ADP-bound NBD, inducing rotation and shifting of subdomain IIB to open the nucleotide-binding cleft and promote ADP release (4, 17). The release of ADP and subsequent binding of ATP trigger the transition of DnaK from the ADP-bound, high substrate affinity state to the ATP-bound, low substrate affinity state. Since NEFs promote the formation of the low substrate affinity state, it is not surprising that they have an influence on the affinity of DnaK for its substrate, as has now been reported by several laboratories (5–9).

While this previous research supports the role of NEFs in the dissociation of substrates from DnaK (and by extension other Hsp70s), the mechanism by which GrpE binding to the DnaK NBD affects substrate affinity was not clear. There has been support, largely based on studies of N-terminally truncated GrpE, that the N-terminal disordered tails of GrpE are key to its facilitation of peptide release (5–9), but no interaction of the tails and the SBD has been observed to date. Additionally, many of these previous studies utilized ADP-bound DnaK, but we have recently shown that the ADP-DnaK/GrpE ternary complex is weak and populates an ensemble of interconverting conformations, with DnaK adopting a more stable GrpE complex in the absence of nucleotide (17).

Published structures of complexes between GrpE and DnaK of *E. coli* homologues have shed light on the nature of the interaction and how it might enable GrpE to modulate substrate affinity. Notably, the *M. tuberculosis* DnaK/GrpE complex (8), the *H. sapiens* Mortalin/GrpEL1 complex (18), and the *G. kaustophilus* DnaK/GrpE complex (19), all revealed a conserved interaction interface between the DnaK SBD and the coiled-coil domain of GrpE, in addition to the previously identified interaction interfaces between the DnaK NBD and GrpE (7, 17). In addition, the structural ensemble of the *M. tubuculosis* DnaK/GrpE complex included some examples with greater proximity of the N-terminal tails to the SBD than observed in many other structures (8). We used the powerful new structure prediction method AlphaFold2 and found that the complex of *E. coli* DnaK and GrpE is very similar to that of the homologues. The conservation of key interaction regions between DnaK and GrpE suggests an evolutionary selection for function, indicating that these interactions are critical to the functionality of the DnaK/GrpE system. Our sequence alignments supported this conservation and thus the importance of the interaction interfaces. It is also reasonable to propose that the interaction of the DnaK NBD with GrpE’s globular domain acts as a driving force for the interaction between the SBD with the coiled-coil domain of GrpE.

We have made a number of observations, largely based on NMR, that taken together support a model in which formation of the DnaK/GrpE complex leads to transient binding of the GrpE N-terminal tails through their conserved ^17^IIM^19^ motif to the canonical substrate-binding site of the DnaK SBD. The important functional role played by weak, transient binding of the GrpE N-terminal tails to the DnaK SBDs is critically reliant on enhanced avidity from local concentration conferred by the formation of the complex between DnaK and GrpE. Consistent with this interpretation is the observation that a peptide consisting of residues 1-33 from GrpE’s N-terminal tails fails to compete with DnaK bound to the model peptide FITC-p5, even when titrated up to 200 μM (**Figure S4**). Therefore, we conclude that the transient, weak binding of the GrpE N-terminal disordered tails to the DnaK SBD are key for *a functional* complex, as GrpE^33-197^ is not able to stimulate peptide release and has been previously shown not to support DnaK assistance of luciferase refolding (5).

The role of GrpE as a thermosensor is also elucidated by our studies of the temperature dependence of the N-terminal tail interaction with the DnaK SBD and its modulation of substrate binding to the DnaK SBD (14, 15). As the temperature increases, the coiled-coil domain of GrpE starts to unfold (13, 16), and GrpE loses its ability to assist DnaK in refolding substrates (9, 14, 15, 20). We conclude from our observations that this loss of GrpE coiled-coil domain stability is accompanied by a loss of the ability to stimulate peptide dissociation and moreover, that binding GrpE’s N-terminal tails to the DnaK SBDs is decreased because of the greater dynamics of a larger segment of the GrpE N-terminal region accounts.

Overall, it is striking how Nature has shaped the interaction between DnaK and GrpE to support efficient protein folding within the cell in a temperature-dependent manner. In addition, the work described provides a compelling example of a system in which structural properties of a complex can enhance the functional importance of a dynamic region of a protein by favoring weak, transient interactions that would otherwise be unlikely to be influential.

## Experimental Procedures

### Protein expression and purification

Wild type *E. coli* DnaK was expressed and purified as described previously (17, 28, 29). *E. coli* GrpE and GrpE^33-197^ were expressed in *E. coli* BL21(*DE3*) and GrpE ASA was expressed in *E. coli* C43(*DE3*) at 37 °C and purified as described previously (17). GrpE concentrations throughout the article always refer to the dimer (17).

To prepare ^13^C methyl label ILV nucleotide free DnaK and ^13^C methyl label ILV nucleotide free NBD^1-392^, established protocols were followed (30–33).

### Sequence alignments

For the DnaK alignments, the following DnaK sequences were used: *Mycobacterium tuberculosis* DnaK (UNIPROT: P9WMJ9), *Escherichia coli* DnaK (UNIPROT: P0A6Y8), *Homo sapiens* Mortalin HSPA9 (UNIPROT: P38646), and *Geobacillus kaustophilus* DnaK (UNIPROT: Q5KWZ7). For the GrpE alignments, the following GrpE sequences were used: *Mycobacterium tuberculosis* GrpE (UNIPROT: P9WMT4), *Legionella pneumoniae* GrpE (UNIPROT: 032481), *Staphylococcus aureus* GrpE (UNIPROT: P63191), *Klebsiella pneumoniae* GrpE (UNIPROT: B5XVJ9), *Enterobacter cloacae* GrpE (UNIPROT: A0A6S5K1M6), *Citrobacter amalonaticus* GrpE (UNIPROT: AOA24RQH2), *Escherichia coli* GrpE (UNIPROT: P09372), *Salmonella enteritidis* GrpE (UNIPROT: B5QUG9), *Clostridioides difficile* GrpE (UNIPROT: Q182F1), *Geobacillus kaustophilus* GrpE (UNIPROT: Q5KWZ6), and *Bacillus subtilis* GrpE (UNIPROT: P15874). Sequence alignments were created using the Clustal Omega (34) and Blast (35, 36) algorithms.

### NMR spectroscopy and chemical-shift analysis

Methyl ^13^C-HMQC spectra were obtained at 25 or 38 °C on a Bruker Avance III 600 MHz instrument and analyzed as described previously (17, 31). ^15^N-HSQC spectra for GrpE, GrpE^33-197^, GrpE/DnaK, GrpE/NBD, and GrpE/DnaK/peptide were acquired at 25 °C using a 600 MHz Bruker Avance Neo spectrometer. Heteronuclear NOEs were measured using an interleaved pulse sequence (Bruker pulseprogram hsqcoef3gpsi) and calculated as the ratio of saturated over non-saturated peak intensity. Resonance assignments for the N-terminal amide signals were obtained using a combination of HNCACB, HNCO, and HN(CA)CO triple-resonance experiments, collected with a 700 MHz Bruker Avance Neo spectrometer.

### Substrate binding

The apparent affinities of DnaK for fluorescein isothiocyanate (FITC)-labeled peptide p5 (ALLLSAPRR) in the absence or presence of GrpE, GrpE^33-197^, or GrpE ASA was measured at 22 or 38 °C in a Biotek Synergy 2 micro plate reader (Biotek), with excitation at 485 nm and emission at 516 nm. 0 - 25 µM DnaK was added to 35 nM FITC-p5 in HMK buffer with 1 mM DTT in triplicate in the absence or presence of 1 µM GrpE, GrpE^33-197^, or GrpE ASA. The mixture was then incubated in 384-well plates for 4 h at 22 or 38 °C, after which the fluorescence anisotropy was measured. The data were analyzed as described previously (27). Statistical analysis was performed using a one-way ANOVA to determine the significance of differences between the GrpE variants. The same process was repeated with varying concentrations of GrpE. Here, 0 - 25 µM GrpE or GrpE^33-197^ was added to 35 nM FITC-p5 in HMK buffer with 1 mM DTT in triplicate in the presence of 0.5 µM DnaK. The same process was also repeated with varying concentrations of a peptide consisting of residues 1-33 from GrpE’s N-terminal tails. Here, 0 - 200 µM GrpE^1-33^ was added to 35 nM FITC-p5 in HMK buffer with 1 mM DTT in triplicate in the presence of 0.5 µM DnaK.

## Data availability

All data are included in this article or in the Supporting Information, except for the structure coordinates predicted by AlphaFold2, which are being deposited in ModelArchive. Resonance assignments for the *E. coli* GrpE disordered tails have been deposited in the BMRB. The latter information can be shared upon request to Akshitha Maqtedar or Lila M. Gierasch.

## Supporting Information

This article contains supporting information.

## Acknowledgments

We thank Jason Gestwicki (University of California San Francisco, USA) for his generous gift of the pMCSgrpe plasmid. We acknowledge the Biophysical Characterization Core Facility (RRID:SCR_019063) and the NMR Core Facility of the Institute for Applied Life Sciences at the University of Massachusetts Amherst for access to excellent instrumentation. Additional NMR data were collected at the Gregory P. Mullen NMR Structural Biology Facility at UConn Health, which is a member of the NSF Network for Advanced NMR.

## Author contributions

L. M. G. conceptualization; A. M., M.-A. R., R. V. M. and E. M. C. methodology; A. M., M.-A. R., and R. V. M. investigation; A. M., M.-A. R., and R. V. M. formal analysis; A. M., M.-A. R., E. M. C, and L. M. G. writing–original draft; A. M., M.-A. R., E. M. C., R. V.M., and L. M. G. writing–review & editing; E. M. C. and L. M. G. visualization; L. M. G. supervision; L. M. G. funding acquisition.

## Funding and additional information

This work was supported by the NIH grant GM18161 to Lila M. Gierasch and NIH grant F32-GM156016 to Robert V. Williams. The NMR Facility at UConn Health as part of the NSF Network for Advanced NMR is supported by NSF grants 1946970 and 2529058. The 800 MHz and 700 MHz instruments at UConn Health were funded by NIH grants S10-RR023041 and S10-OD034297, respectively. The content is solely the responsibility of the authors and does not necessarily represent the official views of the National Institutes of Health.

## Abbreviations

Apo-DnaK: nucleotide-free DnaK
Bag: BAG family molecular chaperone
DnaK: 70-kDa heat shock proteins of prokaryotes
FITC: fluorescein isothiocyanate
GrpE: nucleotide exchange factor of prokaryotes
HMQC: heteronuclear multiple quantum correlation
Hsp70: 70-kDa heat shock proteins
Hsp110: 105-kDa heat shock proteins
HspBP-1: Hsp70-binding protein 1
HSQC: heteronuclear single quantum correlation
app-K_D_: apparent dissociation constant
NBD: nucleotide-binding domain
NEF: nucleotide exchange factor
PDB: protein data bank
SBD: substrate-binding domain

