## Supplemental Information for "The nucleotide exchange factor, GrpE, modulates substrate affinity by interaction of its N-terminal tails with the DnaK substrate-binding domain"

**Contains:**

Supporting figures S1 – S5

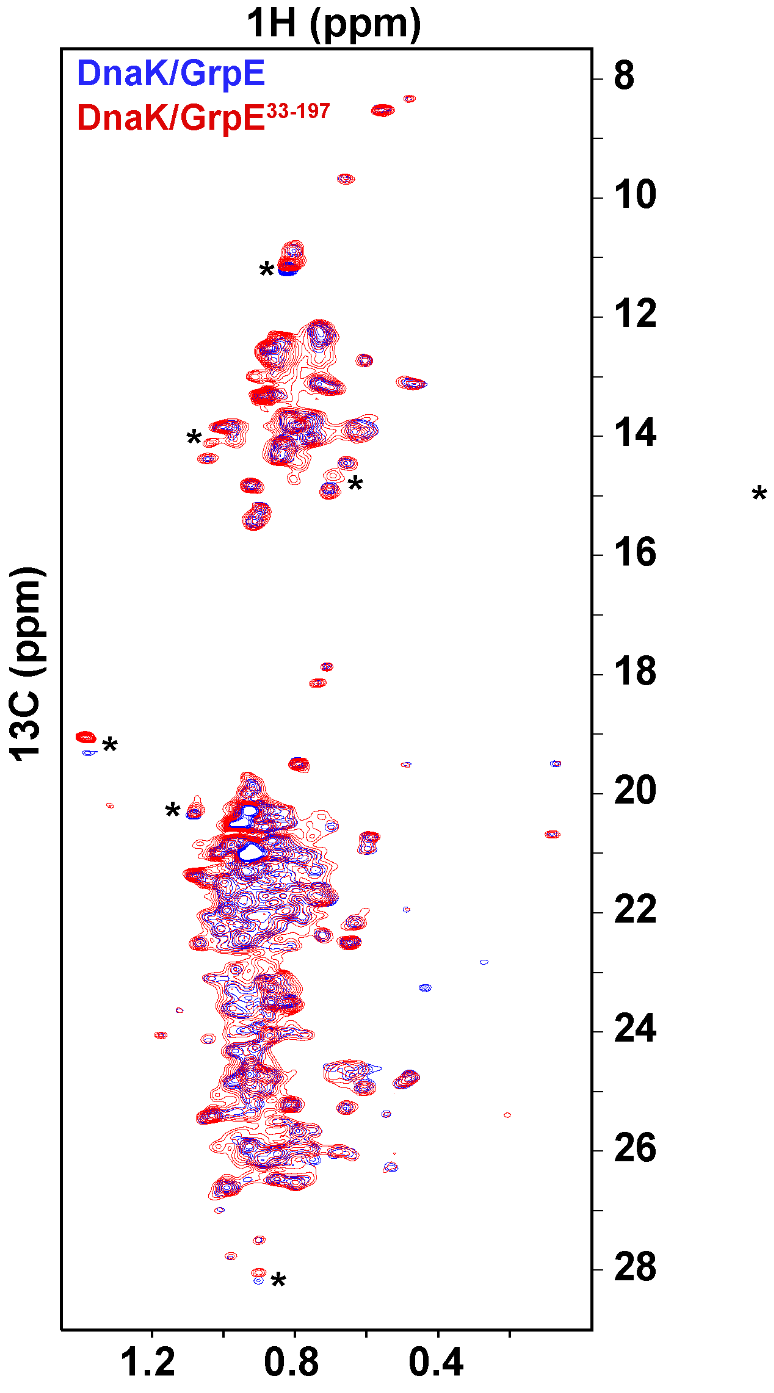


**Figure S1. NMR chemical shift perturbations support binding of GrpE tail to DnaK SBD.**

HMQC of $^{13}$C-methyl labeled ILV-DnaK/GrpE complex (blue) and $^{13}$C-methyl labeled ILV-DnaK/GrpE$^{33\text{-}197}$ complex (red) at 25 °C. Asterisks show residue examples with clear chemical shift perturbations.

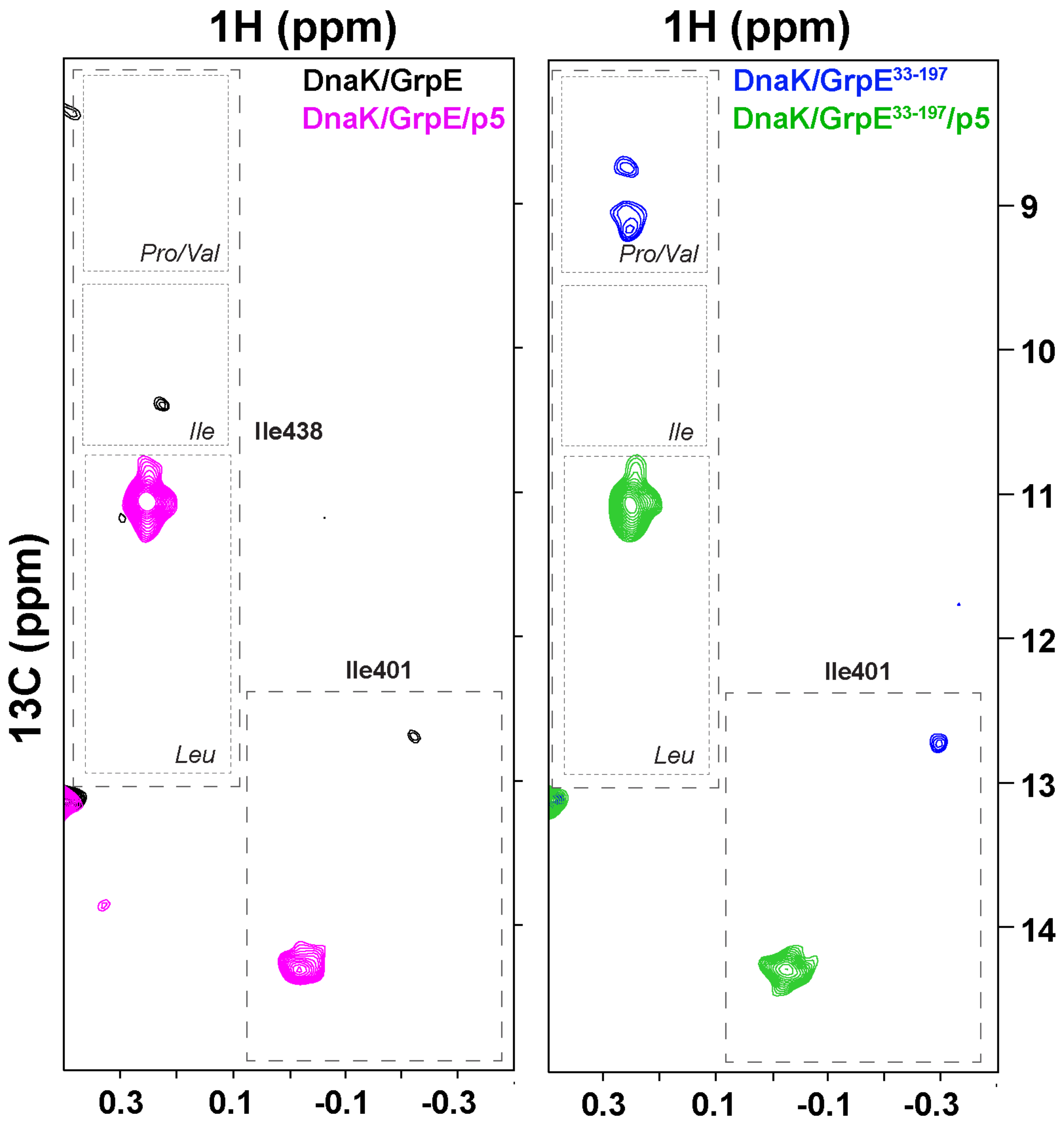


**Figure S2. DnaK SBD reporter methyl NMR signals from residues I401 and I438 indicate the identity of residues binding to the central pocket of the SBD binding groove upon complex formation with GrpE and its variants and when the substrate p5 is added.**

Ile resonances of the substrate binding pocket on the HMQC at 25 °C of (Left) $^{13}C$-methyl labeled ILV-DnaK/GrpE complex (black) and $^{13}C$-methyl labeled ILV-DnaK/GrpE/p5 complex (pink) and (Right) $^{13}C$-methyl labeled ILV-DnaK/GrpE$^{33-197}$ complex (blue) and ILV-DnaK/GrpE$^{33-197}$/ p5 complex (green).

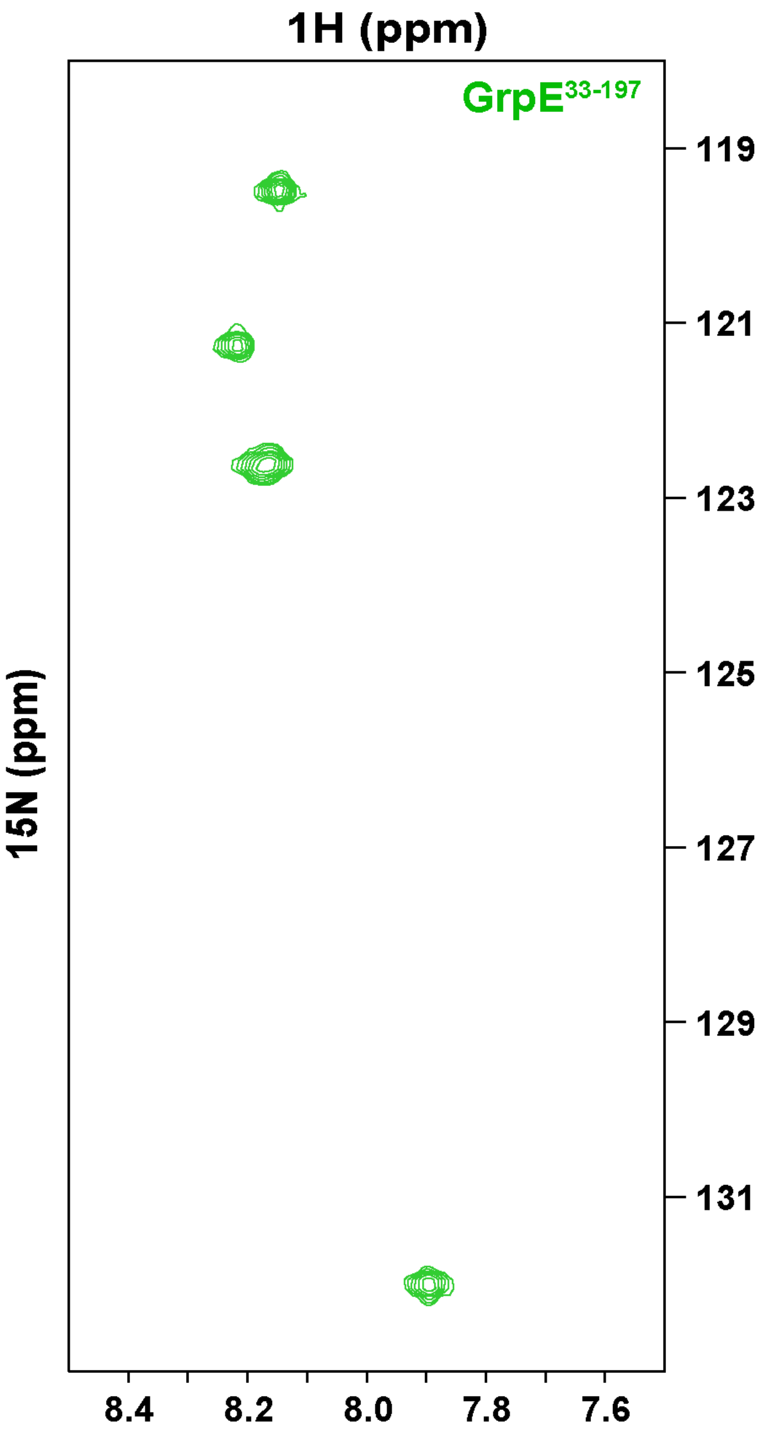


**Figure S3. $^{15}$N-GrpE$^{33-197}$ NMR experiments show that some GrpE residues are dynamic.**

$^{1}$H-$^{15}$N HSQC spectrum of the $^{15}$N -labeled GrpE$^{33-197}$ spectrum (green). There are no assignments for these resonances but we speculate that they arise from the N-terminal residues.

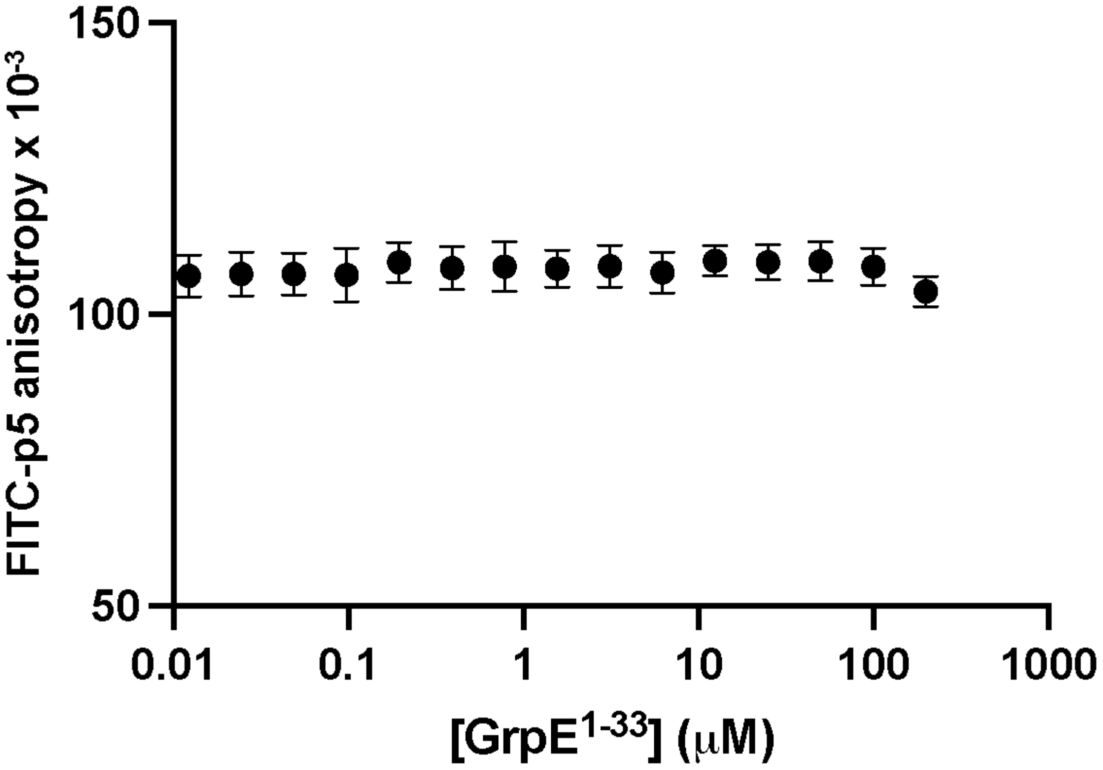


**Figure S4. Peptide consisting of residues 1-33 of GrpE's N-terminal tails fails to compete with the model peptide FITC-p5 for DnaK binding.**

Apparent affinity of DnaK for the model peptide FITC-p5 in the presence of 0-200 μM of a competing peptide consisting of residues 1-33 from GrpE N-terminal tails at 22 °C.

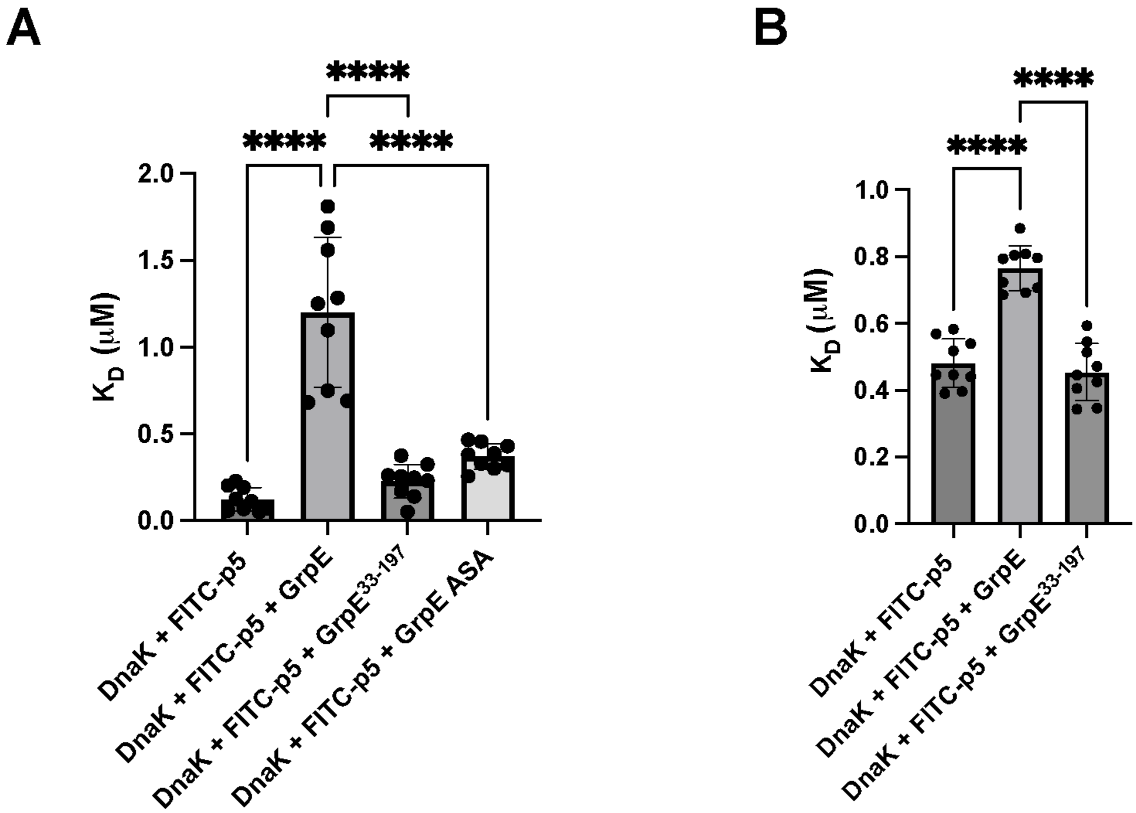


**Figure S5. Statistical analysis for the apparent $K_D$ of DnaK for FITC-p5 in the absence or presence of GrpE and its variants.**

*A,* One-way ANOVA for the apparent $K_D$ of DnaK for the model peptide FITC-p5 in the absence or presence of 1 μM GrpE, $GrpE^{33\text{-}197}$, and GrpE ASA at 22 °C. *B,* One-way ANOVA for the apparent affinity of DnaK for the model peptide FITC-p5 in the absence or presence of 1 μM GrpE and $GrpE^{33\text{-}197}$ at 38 °C.
